# Connectivity between supplementary motor complex and primary motor cortex: a dual-coil paired-pulse TMS study

**DOI:** 10.1101/2024.08.31.610643

**Authors:** Hakjoo Kim, Yuming Lei, Shancheng Bao, Angelina T. Huynh, John J. Buchanan, Jessica A. Bernard, Joshua C. Brown, David L. Wright

## Abstract

In recent years, dual-coil paired-pulse transcranial magnetic stimulation (ppTMS) has garnered interest for its potential in elucidating neural circuit dynamics. In this study, the dual-coil ppTMS was utilized to assess the effective connectivity between the supplementary motor complex (SMC) and the primary motor cortex (M1) in humans. A robust facilitatory connection between the SMC and M1 was observed, manifested as a 19% increase in mean peak-to-peak motor-evoked potentials following preconditioning of SMC 7 ms prior to M1 stimulation. Importantly, the facilitatory influence of SMC only occurred when the preconditioning stimulation was administered 4 cm anterior to Cz but not when applied at 5-cm, 6-cm, or 7-cm distance. While previous work has focused on demonstrating important temporal dynamics for SMC-M1 plasticity, the present findings highlight a critical contribution of spatial specificity for modulation of SMC-M1 circuitry.

**Graphical abstract:** 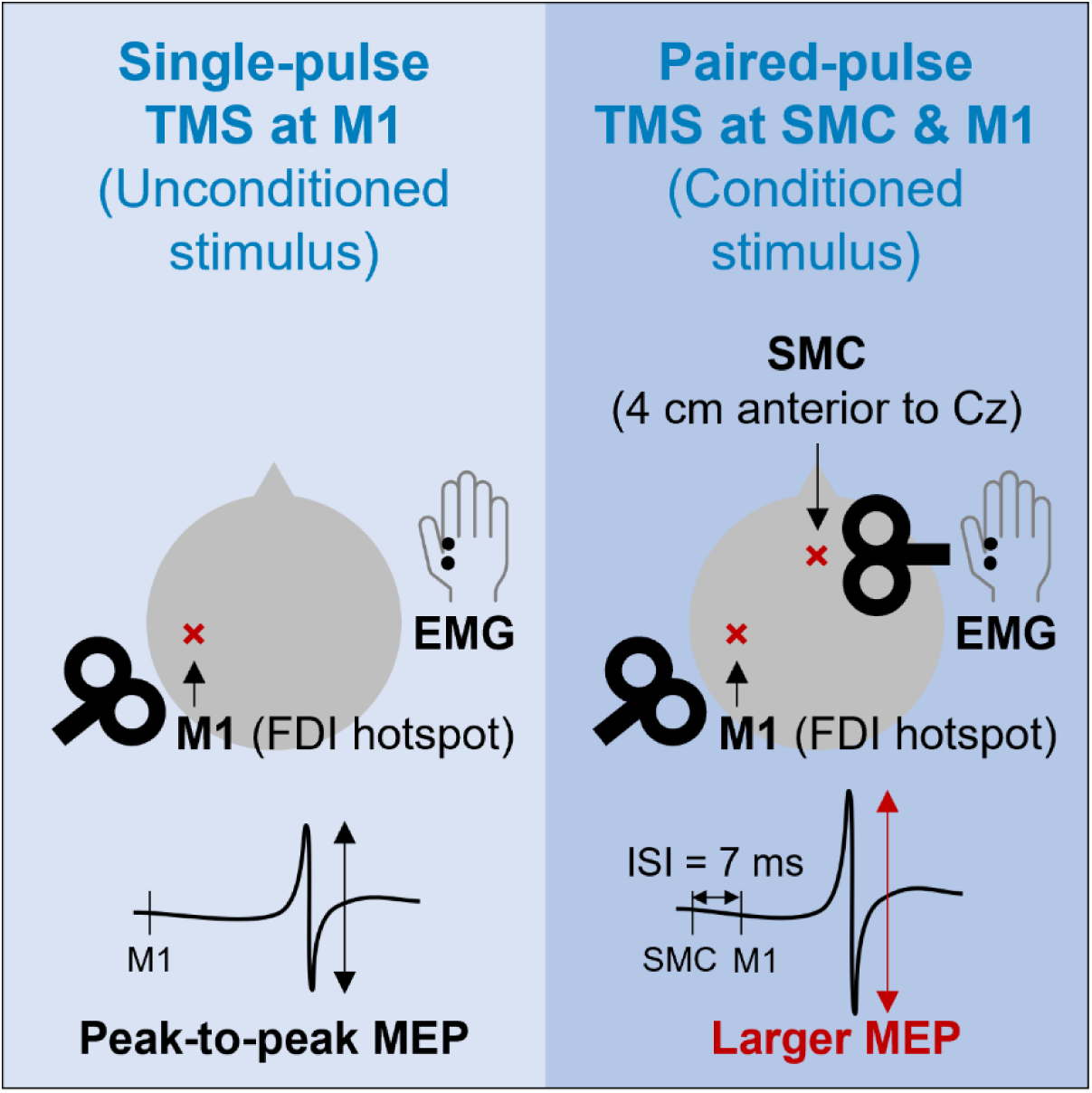

**Highlights:**

- The conditioned stimulus induced larger MEPs compared to the unconditioned stimulus
- The connectivity between SMC and M1 diminished when the SMC coil was moved forward
- Findings highlight the importance of spatial specificity for modulation of SMC-M1 circuitry

## INTRODUCTION

The supplementary motor complex (SMC) is part of the superior frontal gyrus, specifically located on the medial wall of the brain, labeled Brodmann’s area 6. For humans, this area lies anterior to the leg representation of the primary motor cortex (M1) (Nachev et al., 2008; Cona & Semenza, 2017). Functionally, SMC has been implicated in the performance of sequential movements (Tanji, 2001; Nachev et al., 2008). In the case of primates, findings from studies using single-unit recordings (Clower & Alexander, 1998; Shima & Tanji, 2000) and through the administration of pharmacological agents (Shima & Tanji, 1998) have highlighted the selective involvement of cells within SMC when planning and initiating motor sequences. In humans, early imaging work by Roland et al. (1980) revealed heightened regional cerebral blood flow at SMC prior to executing a sequence of ballistic finger movements (see also Grafton et al.,1995; Wiestler & Diedrichsen, 2013). A central role of SMC for preparing action sequences has been verified using transcranial magnetic stimulation (TMS). For example, Gerloff et al. (1997) revealed disrupted performance of complex finger sequences by administering 15-Hz repetitive TMS to SMC but not to other motor and parietal sites (also see Verwey et al., 2002; Kennerley et al., 2004). During the early phase of skill learning that involves novel sequential content, the more rostral portion of the SMC, referred to as the pre-supplementary motor area (SMA), has been identified as particularly critical, whereas the SMA proper, the more caudal segment of this region, appears more crucial for the execution of well-learned actions (Hikosaka et al., 1996; Sakai et al., 1998; Sakai et al., 1999).

Early understanding of SMC involvement in both the control and learning of complex sequential behaviors benefited from the extensive development in neuroimaging techniques during the last few decades (Kim et al., 2010). One consequence of such advancements is the identification of intra-and inter-regional connectivity patterns that are associated with enhanced acquisition and retention of complex sequential actions. Unraveling causal relationships, however, has relied on the use of non-invasive brain stimulation tools such as TMS to uncover the direct and indirect influence of one neural site (e.g., SMC) on another (i.e., M1), referred to as effective connectivity (Derosiere & Duque, 2020; Neige et al., 2021).

During the last twenty years, dual-coil paired-pulse TMS (ppTMS) has been used with humans to determine if a facilitatory or inhibitory tone is exerted by one region on another. For example, ppTMS has been used quite extensively to probe the influence of the cerebellum on M1, or, in other words, cerebellar-M1 connectivity (van Malderen et al., 2023). Early work by Ugawa and colleagues (1995) demonstrated that administering a single TMS pulse at the cerebellum 5 ms before a second separate pulse was delivered at M1 led to a down-regulation in the excitability at M1, manifested as a reduction in the amplitude of motor-evoked potential (MEP) when compared to a condition that involved stimulation of M1 only. Since these initial efforts, ppTMS has been employed to detail a host of inter- and intra-hemispheric interactions involving M1 and a variety of neural regions that make significant contributions to the performance of action sequences (see van Malderen et al., 2023 for a detailed review of this work).

Despite the general acceptance that SMC-M1 connectivity has functional relevance, there are only a modest number of studies that have adopted ppTMS to probe this part of the cortico-motor network. To date, ppTMS studies that were designed to examine the influence of SMC on M1 have mostly focused on detailing the temporal dynamics of this circuitry. For example, Arai et al. (2011) offered preliminary evidence for timing-dependent plasticity of the SMC-M1 cortical network that persisted for up to 30 minutes. Specifically, they demonstrated increased MEP amplitude when suprathreshold stimulation of SMC preceded M1 stimulation by 6 ms but not 15 ms. Subsequent work extended the findings of Arai and colleagues, revealing that SMC’s facilitatory influence on M1 at an inter-stimulus interval (ISI) of 6 ms was more pronounced for younger as opposed to older adults. Recently, Rurak et al. (2021) revealed that an ISI of 7 ms resulted in the most reliable facilitatory impact of SMC on M1 for both young and older individuals. Importantly, the magnitude of facilitation impacted by SMC on M1 appears to be functionally relevant as it was positively correlated with bimanual performance (Green et al., 2018).

In the aforementioned studies, the examination of the temporal dynamics of SMC-M1 connectivity generally adopted a distance of ∼4 cm anterior to Cz in the international 10-20 system as the appropriate anatomical location of SMC for the administration of exogenous stimulation on the surface of the skull used as a part of the ppTMS protocol (Arai et al., 2011; Arai et al., 2012; Lu et al., 2012). In two studies (Arai et al., 2011; Green et al., 2018), the conditioning stimulation at SMC during ppTMS was administered at ∼7 cm anterior to Cz to probe potential pre-SMA modulation of M1. In both cases, the anticipated change in the MEP amplitude when compared to M1 stimulation alone was absent, suggesting topographic specificity for the facilitatory effect induced by SMC preconditioning. The present work extended the investigation of the spatial specificity of SMC modulatory influence on M1. Specifically, the distance from Cz at which the conditioning stimulation was administered prior to M1 stimulation was systematically manipulated from 4 cm to 7 cm anterior to Cz in 1-cm steps. For all conditions, an ISI of 7 ms was used between the conditioning and test stimuli (see Rurak et al., 2021). First, congruent with previous findings (Arai et al., 2012; Green et al., 2018; Rurak et al., 2021), it was expected that the MEP amplitude following preconditioning at 4 cm would be increased, verifying the facilitatory tone SMC exerts on M1. Second, it was anticipated that any modulation of M1 excitability via preconditioning of SMC when administered at 5, 6, or 7 cm would exert a smaller impact on the resultant MEP amplitude observed at M1 (see Arai et al., 2011; Green et al., 2018).

## RESULTS

### Administering a preconditioning stimulus at 4 cm anterior to Cz increases M1 excitability

The initial issue addressed involved verifying that preconditioning SMC at 4 cm anterior to Cz could lead to an elevation in M1 excitability. Figure 1 displays individual and mean peak-to-peak MEPs when TMS was applied at M1 only (unconditioned stimulus, US) and at both SMC and M1 (conditioned stimulus, CS), where SMC stimulation was administered at 4 cm anterior to Cz (*n* = 21) in a manner similar to that adopted in previous studies addressing SMC-M1 connectivity (Arai et al., 2011; Green et al., 2018; Rurak et al., 2021). A paired samples *t*-test was used to compare the mean peak-to-peak MEP amplitude for the 21 individuals who experienced the US and CS conditions at 4 cm anterior to Cz. However, this comparison was not made for the 42 individuals who received SMC stimulation at 5, 6, or 7 cm anterior to Cz (5 cm: *n* = 20; 6 cm: *n* = 15; 7 cm: *n* = 7). The mean peak-to-peak MEP amplitude for the US condition (*M* = 0.539 mV, *SEM* = 0.077 mV) was significantly smaller than that observed in the CS condition (*M* = 0.643 mV, *SEM* = 0.091 mV), *t*(20) = -3.229, *p* = .004. The mean peak-to-peak MEP amplitude for the CS condition was 19.33% larger than for the US condition. Of the 21 individuals assessed, the peak-to-peak MEP amplitudes were increased for ∼76% of participants when M1 was preconditioned by SMC stimulation, suggesting a robust influence of SMC on M1.

**Figure 1.**
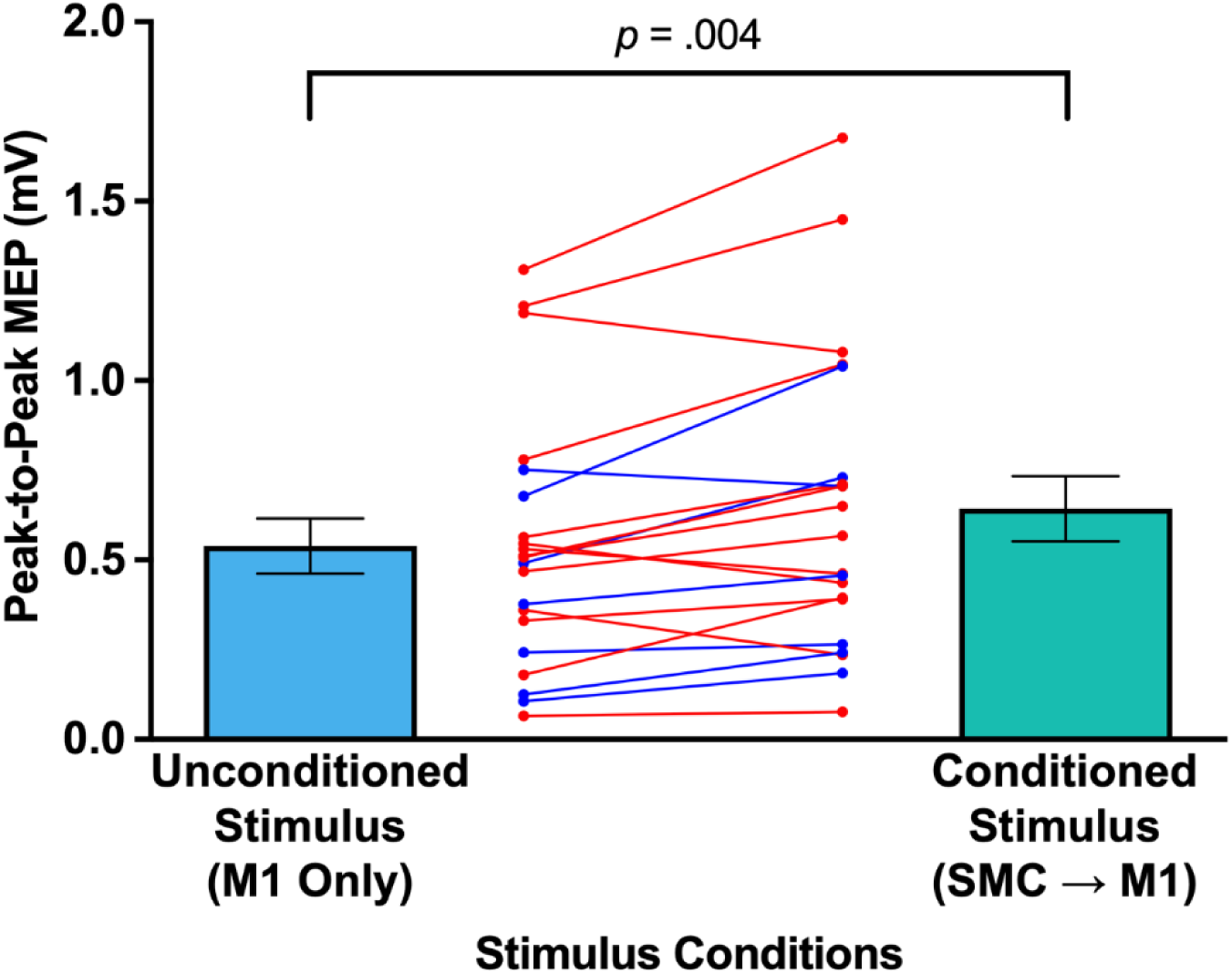
Peak-to-Peak MEPs when TMS applied at M1 only and both SMC and M1. The left bar (blue) denotes unconditioned stimulus (i.e., single-pulse TMS at M1 only), and the right bar (mint) denotes conditioned stimulus (i.e., ppTMS at SMC prior to M1 stimulation). The red lines correspond to female participants and the blue lines to males. The error bars represent the standard errors.

### Administering preconditioning stimulus beyond 4 cm anterior of Cz eliminates facilitation of M1 excitability

A second issue addressed herein was the impact of administering the preconditioning stimulus in the CS condition more anterior to the standard location of 4 cm anterior of Cz used in previous work on the resultant M1 excitability. To examine this issue, a CS to US MEP ratio was determined for each individual for each of the different locations at which the preconditioning stimulus was applied in the CS condition. In this analysis, CS to US MEP ratios of the preconditioning stimulus that was applied at 4 cm, 5 cm, 6 cm, or 7 cm anterior to Cz were analyzed. A CS to US ratio > 1 indicates facilitation in M1 excitability, whereas a ratio of < 1 reveals a reduction in M1 excitability as a result of preconditioning. Figure 2 displays both individual and mean CS to US ratios as a function of the location of the preconditioning stimulus relative to Cz.

**Figure 2.**
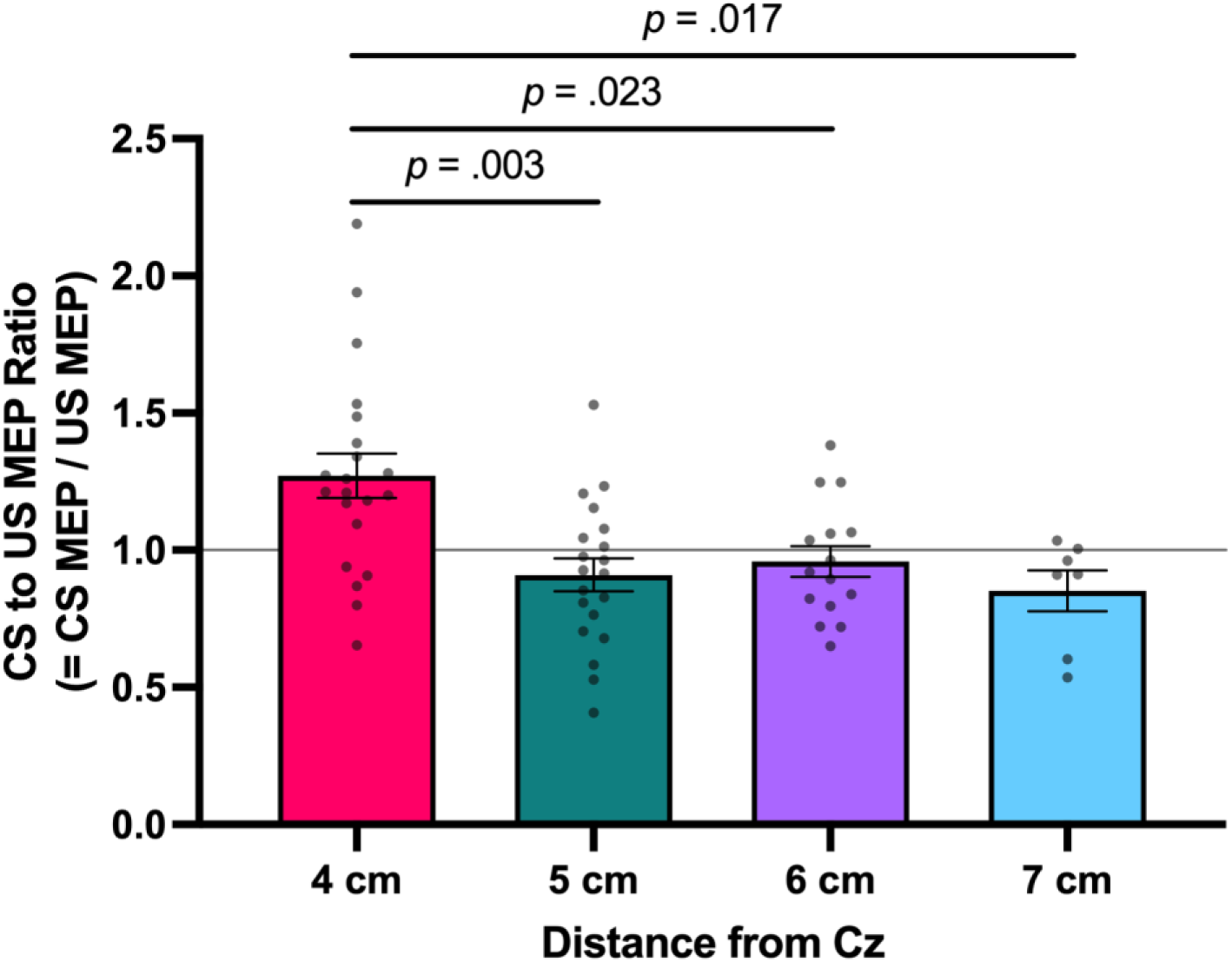
Conditioned stimulus (CS) to unconditioned stimulus (US) MEP ratios as a function of location of preconditioning stimulus relative to Cz. From the left, bars denote the CS to US ratios when SMC stimulation was administered at 4, 5, 6, or 7 cm anterior to Cz. Values greater than 1 indicate facilitation in MEP amplitude as a result of preconditioning. The error bars represent the standard errors.

It should be noted that participants were only exposed to one location for preconditioning and that the 63 participants in the experiment were not evenly distributed across the preconditioning locations (4 cm: *n* = 21; 5 cm: *n* = 20; 6 cm: *n* = 15; 7 cm: *n* = 7). The CS to US MEP ratio for each individual for each preconditioning location was submitted to a one-way between-subject (location of preconditioning) analysis of variance (ANOVA). Levene’s test confirmed the homogeneity of variance across the four locations of preconditioning (*p* = .443). The one-way between factor ANOVA revealed a statistical difference in the mean CS to US MEP ratio as a function of location of preconditioning, *F*(3, 59) = 7.172, *p* < .001, partial *η*^2^ = 0.267. Subsequent Scheffe post hoc testing revealed that the mean CS to US MEP ratio at the 4-cm location (*M* = 1.271, *SEM* = 0.081) was statistically larger than that observed at the 5-cm (*M* = 0.910, *SEM* = 0.059, *p* = .003), 6-cm (*M* = 0.958, *SEM* = 0.056, *p* = .023), and 7-cm locations (*M* = 0.852, *SEM* = 0.075, *p* = .017) (see Figure 2). In addition, the differences in CS to US MEP ratios between the 5, 6, and 7-cm locations did not differ significantly.

## DISCUSSION

### Influence of SMC conditioning at 4 cm anterior to Cz on M1

An important objective of this study was to confirm the facilitatory influence of SMC conditioning at 4 cm anterior to Cz on M1, as reported in the previous studies (Arai et al., 2012; Green et al., 2018; Rurak et al., 2021). Specifically, Rurak and colleagues demonstrated that administering a conditioning stimulus at 4 cm anterior to Cz, 7 ms before M1 stimulation, led to an approximate 20% increase in the peak-to-peak MEP amplitude. This facilitation of the output from M1, when preceded by SMC stimulation, has been thought to be due to the activation of glutamatergic excitatory interactions between these neural sites (Muakkassa & Strick, 1979; Luppino et al., 1993; Shima & Tanji, 1998).

In the current study, a ppTMS protocol similar to that used by Rurak et al. (2021) was employed, and as expected, the results were consistent with those previously reported (Arai et al., 2012; Green et al., 2018; Rurak et al., 2021). Indeed, the overall facilitatory influence of SMC conditioning on M1 was comparable to that observed by Rurak and colleagues, showing an approximately 20% increase in peak-to-peak MEP amplitude (see Figures 1 and 2). It is noteworthy that the facilitatory influence of SMC on M1 observed in this study was robust, with approximately 76% of participants exhibiting this form of SMC-M1 plasticity (see Figure 1). However, the data also suggest that the modulation of M1 activity by SMC via TMS may depend on the individual’s susceptibility to exogenous stimulation, similar to other non-invasive brain stimulation techniques, such as transcranial direct current stimulation (tDCS) (López-Alonso et al., 2014; Willmot et al., 2024). This is reflected in the fact that a small number of individuals exhibited either no effect or actually experienced reduced M1 activity following preconditioning at SMC.

An alternative explanation for our findings and those reported by Rurak et al. (2021) is that the increased output from M1 in the CS condition might result from direct stimulation of M1 rather than being mediated indirectly through input from SMC, the preconditioning site. This would suggest that the CS condition may have elicited intracortical facilitation (ICF). However, it is important to note that ICF is typically induced with an ISI of 10-15 ms (Kujirai et al., 1993) rather than the shorter ISI of 7 ms used in the current study and by Rurak et al. In fact, Rurak and colleagues examined the stability of the facilitatory influence of SMC on M1 across various ISIs, including 6, 7, and 8 ms, revealing that an ISI of 7 ms was the most reliable in both younger and older adults. Therefore, increasing the ISI towards the time frame frequently used to elicit ICF did not result in a change in M1 output in the study by Rurak et al.

Some additional evidence counter to a direct impact of the conditioning stimulus on M1 can be drawn from Arai et al. (2011), who revealed that MEPs increased when SMC stimulation occurred 6 ms before M1 stimulation (i.e., -6 ms) while decreased when SMC stimulation was applied 15 ms after M1 (i.e., +15 ms). Perhaps more importantly, Neige et al. (2023) recently reported no impact on M1 from SMC stimulation 15 ms before M1 stimulation. While it is impossible to rule out the possibility that heightened M1 excitability in the CS condition of the present experiment and others did not result from a direct influence on M1 rather than being modulated via SMC activity, the existing evidence suggests this is unlikely, given the tight temporal dynamics associated with the facilitatory effect of SMC on M1 (Arai et al., 2011; Rurak et al., 2021; Neige et al., 2023).

### The facilitatory influence of SMC conditioning on M1 is eliminated when the preconditioning location is moved more anterior from Cz

To date, attempts to investigate the tonic influence of SMC activation on M1 corticomotor excitability have primarily concentrated on outlining the crucial temporal dynamics of the SMC-M1 network. Notably, robust facilitation at M1 from SMC conditioning was observed with a 7-ms ISI (Rurak et al., 2021). However, using an ISI of 6 or 8 ms has shown significantly greater variability in corticomotor output changes for both older and younger adults (Rurak et al., 2021).

An opportunity to examine the spatial specificity of the SMC-M1 connectivity discussed in the previous section inadvertently emerged as a result of variations in head size among participants or the specific locations of the M1 hotspots. In several cases, the standard 4-cm distance from Cz could not be used because participants’ smaller head sizes did not allow for the required distance between the two 30-mm TMS coils used to independently stimulate SMC and M1. In such cases, SMC stimulation was administered at a distance that allowed for proper placement of the two TMS coils. Consequently, a significant number of individuals used distances of 5, 6, or 7 cm anterior to Cz instead of 4 cm. This situation allowed for an evaluation of the spatial specificity of the facilitatory influence of SMC on M1 by examining the impact on M1 excitability when the conditioning stimulus was applied at these more anterior locations from Cz.

Assuming that the 4-cm site is an appropriate spatial approximation of SMC on the scalp (Arai et al., 2012; Green et al., 2018; Rurak et al., 2021) and that the SMC-M1 connectivity is responsible for the facilitatory influence discussed earlier, one would expect that applying the same conditioning stimulus at more anterior locations would result in a systematic change in corticomotor excitability observed at M1. For instance, the facilitatory influence might diminish as the conditioning site moves further away. Alternatively, it is possible that no facilitatory influence would be observed beyond the 4-cm location, indicating a high degree of spatial specificity for the impact discussed earlier and by others (Arai et al., 2011; Green et al., 2018). Another possibility is that the facilitatory influence of SMC conditioning on M1 might still be observed even at these more anterior sites, suggesting that M1 output is influenced by circuits extending beyond the 4-cm region anterior to Cz.

As shown in Figure 2, the CS condition involving all sites beyond 4 cm failed to increase M1 excitability. Moving the stimulation site systematically anterior to Cz beyond the 4-cm location eliminated the facilitatory influence of SMC on M1 (cf. Arai et al., 2011; Green et al., 2018). This suggests that the previously reported upregulation of M1 activity is quite focal, limited to a circuit influencing M1 from cells localized to a specific spatial location along the midline, approximately 4 cm anterior to Cz, which Rurak et al. (2021) and others (Arai et al., 2012; Green et al., 2018) attribute to the SMC. Thus, it appears that cells within SMC, located at 4 cm anterior to Cz, directly influence M1, exerting a facilitatory effect that leads to a substantial increase in M1 output, as reflected in larger peak-to-peak MEPs. The upregulation of M1 excitability by SMC input seems to be highly specific in both spatial and temporal domains. While temporal specificity has been suggested in previous studies (see Arai et al., 2011; Rurak et al., 2021), the present study underscores the spatial specificity of SMC’s influence on M1.

## STAR★METHODS

Detailed methods are provided in the online version of this paper and include the following:

- KEY RESOURCES TABLE
- RESOURCE AVAILABILITY

○ Lead contact
○ Materials availability
○ Data and code availability
- MODEL AND STUDY PARTICIPANT DETAILS
- METHOD DETAILS
- AND STATISTICAL ANALYSIS

## ACKNOWLEDGMENTS

Funds from the Omar Smith Endowed Chair in Kinesiology awarded to D.L.W. and from THE SYDNEY AND J.L. HUFFINES INSTITUTE FOR SPORTS MEDICINE AND HUMAN PERFORMANCE awarded to H.K. supported this study.

We would like to extend our sincere gratitude to Dr. James Fluckey for graciously allowing the use of laboratory resources, which significantly contributed to the success of this research.

## AUTHOR CONTRIBUTIONS

Conceptualization, H.K., Y.L., and D.L.W.; Methodology, H.K., Y.L., S.B., and D.L.W.; Software, H.K., Y.L., S.B., and D.L.W.; Validation, H.K., Y.L., S.B., A.H., J.J.B., J.A.B., and D.L.W.; Formal Analysis, H.K., Y.L., J.J.B., J.A.B, and D.L.W.; Investigation, H.K. and A.H.; Resources, H.K., Y.L., S.B., and D.L.W.; Data Curation, H.K., Y.L., J.J.B., J.A.B, and D.L.W.; Writing – Original Draft, H.K., Y.L., J.J.B., J.A.B., J.C.B, and D.L.W.; Writing – Review & Editing, H.K., Y.L., S.B., A.H., J.J.B., J.A.B., J.C.B, and D.L.W.; Visualization, H.K., Y.L., and D.L.W.; Supervision, H.K., Y.L., and D.L.W.; Project Administration, H.K., Y.L., and D.L.W.; Funding Acquisition, H.K. and D.L.W.

## DECLARATION OF INTERESTS

The authors declare no competing interests.

## STAR★METHODS

### KEY RESOURCES TABLE

| REAGENT or RESOURCE | SOURCE | IDENTIFIER |
| --- | --- | --- |
| Software and algorithms |  |  |
| Brainsight (Version 2.4.3) | Rogue Research | https://www.rogue-research.com/tms/brainsight-tms/ |
| Signal (Version 7.04) | CED Ltd. | https://ced.co.uk/products/signal |
| SPSS (Version 28.0.0) | IBM | https://www.ibm.com/spss?lnk=flatitem |
| Prism (Version 10.3.0) | GraphPad Software | https://www.graphpad.com/ |

### RESOURCE AVAILABILITY

#### Lead contact

Further information and requests for resources should be directed to and will be fulfilled by the lead contact, David L. Wright.

#### Materials availability

This study did not generate new unique reagents.

#### Data and code availability

- All data reported in this paper will be shared by the lead contact upon request.
- This paper does not report original code.
- Any additional information required to reanalyze the data reported in this paper is available from the lead contact upon request.

### EXPERIMENTAL MODEL AND STUDY PARTICIPANT DETAILS

#### Participants

63 right-handed undergraduate students (45 females and 18 males, mean age ± SD: 19.78 ± 1.22, age range: 18 to 22) from the Department of Kinesiology and Sport Management at Texas A&M University participated in this study. Prior to participating in this study, all participants provided written informed consent, which was approved by the Texas A&M University Institutional Review Board. Each individual also completed the short version (Veale, 2014) of the Edinburgh Handedness Inventory (Oldfield, 1971) and a prescreening form for TMS. None of the individuals had metallic hardware on their scalp, cardiac pacemakers, implanted medication pumps, intracardiac lines, or central venous catheters. Additionally, they had no history of cortical stroke, other cortical lesions such as brain tumors, seizures, epilepsy, or previous brain surgeries. None had electrical, mechanical, or magnetic implants, nor did they suffer from uncontrolled migraines. Participants were not on prescription medications for brain-related disorders, did not have unstable medical conditions, and had no metal on their body or clothing above the shoulders. None of the individuals were professional musicians. No adverse effects of single-pulse or ppTMS protocol were reported either during or after their participation. Upon completion of the study, participants received course credit for an undergraduate kinesiology class. Table 1 shows the demographic information of the current study.

**Table 1.** Demographic information. SD: standard deviation, LQ: laterality quotient score. *-100 to -61: left handers, -60 to 60: mixed handers, 61 to 100: right handers.

| Condition | *n* (Total) | *n* (Female) | *n* (Male) | Age (SD) | LQ* (SD) |
| --- | --- | --- | --- | --- | --- |
| 4 cm | 21 | 14 | 7 | 19.48 (1.12) | 96.43 (7.01) |
| 5 cm | 20 | 13 | 7 | 19.85 (1.23) | 95.00 (11.75) |
| 6 cm | 15 | 14 | 1 | 19.87 (1.30) | 96.67 (9.99) |
| 7 cm | 7 | 4 | 3 | 20.29 (1.38) | 98.21 (4.72) |
| **Total** | **63** | **45** | **18** | **19.78 (1.22)** | **96.23 (9.16)** |

### METHOD DETAILS

#### Dual-coil paired-pulse transcranial magnetic stimulation (ppTMS)

In order to find a hotspot of the right first dorsal interosseous (FDI) muscle, single-pulse TMS was administered by DuoMAG MP-Dual (DEYMED Diagnostic s.r.o., Czech Republic) with a butterfly T-shaped coil (2 × 30 mm-diameter windings, 30BFT-shaped, DEYMED Diagnostic s.r.o., Czech Republic). An FDI hotspot was determined based on the peak-to-peak MEP amplitudes obtained from pre-set-grid-based hotspot-hunting (grid spacing: 2 by 2 mm, Kim et al., 2023). The hotspot hunting was performed with a TMS coil orientation of a 45-degree angle. After hotspot hunting, the resting motor threshold (rMT) of the FDI muscle was determined as a minimum TMS intensity to evoke the peak-to-peak amplitude of 0.05 mV in at least 5 out of 10 trials (Rossini et al., 2015). Brainsight TMS Navigation system (Version 2.4.3, Rogue Research, Canada) was used to navigate TMS, allowing for consistent stimulation of identical target points. Using the neuronavigation system, when single-pulse TMS and ppTMS were administered at each target, both angular and twist errors were kept to less than 0.05 degrees.

Based on the previous studies (Arai et al., 2012; Green et al., 2018; Rurak et al., 2021), SMC stimulation was administered at 4 cm anterior to the Cz in the international 10-20 system. Additionally, if the hotspot of the FDI muscle was too close to an area 4 cm anterior to Cz, participants received SMC stimulation at 5, 6, or 7 cm anterior to Cz instead of 4 cm (i.e., between-subjects design). The SMC stimulation (i.e., conditioning stimulus) occurred 7 ms prior to M1 stimulation (i.e., test stimulus) at an FDI hotspot (Rurak et al., 2021). Participants received 30 randomly presented unconditioned (i.e., 15 single-pulse TMS at M1, US) and conditioned (i.e., 15 ppTMS at SMC and M1, CS) stimuli (cf. Cavaleri et al., 2017) that were induced by Signal software (Version 7.04, CED Ltd., United Kingdom). The TMS intensity at the left M1 was set to 110% of the rMT (Kallioniemi & Julkunen, 2016). For the SMC, the TMS intensity was set at 140% of the rMT. The coil was positioned at a 45-degree angle to the midline of the brain for M1 stimulation, while for SMC stimulation, the coil was oriented at 270 degrees (Arai et al., 2012; Rurak et al., 2021). The mean peak-to-peak MEPs of the right FDI muscle were compared to quantify the excitability of the corticospinal pathway for both US and CS conditions.

#### Electromyography (EMG)

EMG signals from the right FDI muscle were recorded through disposable Ag-AgCl electrodes (Ambu Neuroline 720 surface electrodes, Ambu A/S, Denmark), amplified by NL844 AC Pre-amplifier (Gain × 100, Digitimer Ltd., United Kingdom), which was connected to NL820A Isolation Amplifier (Gain × 1, Digitimer Ltd., United Kingdom), filtered by NL136 Four Channel Low Pass Filters (2 kHz, Digitimer Ltd., United Kingdom), and sampled at 5 kHz by Signal software (Version 7.04, CED Ltd., United Kingdom). TMS-induced MEPs were recorded from the right FDI muscle, and the background noise was kept to less than 0.02 mV.

#### Procedure

Prior to participation, all individuals had submitted a signed consent form and prerequisite questionnaires. An experimenter measured each participant’s head size with a measuring tape and then located and marked two specific points on the scalp: Cz and a point 4 cm anterior to Cz. If the conditioning TMS coil could not be positioned at the SMC stimulation point (i.e., 4 cm anterior to Cz) due to small head size or the hotspot being too near the conditioning point, alternative spatial locations at 5, 6, and 7 cm anterior to Cz were marked. Participants who could not receive stimulation at the 4-cm point were then given SMC stimulation at the closest possible location to Cz among the 5, 6, or 7-cm point. The conditioning points (i.e., 4, 5, 6, or 7 cm anterior to Cz) were registered in the neuronavigation system by administering a minimum of three single TMS pulses. The final conditioning target was determined by averaging the locations of these pulses (cf. the center of gravity). Single-pulse TMS was used to find a hotspot of the right FDI muscle and determine the rMT. Subsequently, participants received paired-pulse TMS at the conditioning point and the hotspot. The experimental procedure was completed within one hour, including obtaining informed consent.

### QUANTIFICATION AND STATISTICAL ANALYSIS

The primary dependent variable of the present study was the peak-to-peak MEPs of the right FDI muscle. The peak-to-peak MEPs were captured by Signal (Version 7.04, CED Ltd., United Kingdom). From 30 random US and CS presentations, raw MEPs from 15 US and raw MEPs from 15 CS were separated for statistical analysis. Among the 63 participants, individuals who received SMC stimulation at 5, 6, and 7 cm anterior to Cz were analyzed separately. MEP ratio (= CS MEP / US MEP) was calculated to compare the difference between distances of 4, 5, 6, and 7 cm. Statistical analyses, such as paired samples *t*-test and one-way ANOVA (alpha level of 0.05), were conducted using SPSS (Version 28.0.0, IBM, United States), and graphs were generated by Prism (Version 10.3.0, GraphPad Software, United States).

## Notes

### Competing Interest Statement

The authors have declared no competing interest.

